# Epitope - based peptide vaccine against *Fructose-bisphosphate aldolase (FBA)* of *Madurella mycetomatis* using immunoinformatics approaches

**DOI:** 10.1101/352625

**Authors:** Arwa A. Mohammed, Ayman M. H. ALnaby, Solima M. Sabeel, Fagr M. AbdElmarouf, Amina I. Dirar, Mostafa M. Ali, Mustafa A. Khandgawi, Abdelhameed M. Yousif, Eman M. Abdulgadir, Magdi A. Sabahalkhair, Ayman E. Abbas, Mohammed A. Hassan

## Abstract

**Background:** Mycetoma is a distinct flesh eating and destructive neglected tropical disease. It is endemic in many tropical and subtropical countries. Mycetoma is caused by bacterial infections (actinomycetoma) such as Streptomyces somaliensis and Nocardiae or true fungi (eumycetoma) such as Madurella mycetomatis. Until date, treatments fail to cure the infection and the available marketed drugs are expensive and toxic upon prolonged usage. Moreover, no vaccine was prepared yet against mycetoma.

**The aim** of this study is to predict effective epitope-based vaccine against fructose-bisphosphate aldolase enzymes of M. mycetomatis using immunoinformatics approaches.

**Methods and Materials:** Fructose-bisphosphate aldolase of *Madurella mycetomatis* Sequence was retrieved from NCBI. Different prediction tools were used to analyze the nominee’s epitopes in Immune Epitope Database for B-cell, T-cell MHC class II & I. Then the proposed peptides were docked using Autodock 4.0 software program.

**Results and Conclusions:** The proposed and promising peptides KYLQ shows a potent binding affinity to B-cell, FEYARKHAF with a very strong binding affinity to MHC1 alleles and FFKEHGVPL that show a very strong binding affinity to MHC11and MHC1 alleles. This indicates a strong potential to formulate a new vaccine, especially with the peptide FFKEHGVPL which is likely to be the first proposed epitope-based vaccine against Fructose-bisphosphate aldolase of Madurella mycetomatis. This study recommends an in-vivo assessment for the most promising peptides especially FFKEHGVPL.

## 1. Introduction

Mycetoma is a distinct flesh eating and a devastating neglected tropical disease, caused by bacterial infections (actinomycetoma) such as *Streptomyces somaliensis* and Nocardiae or true fungi (eumycetoma) such as *Madurella mycetomatis* ^[1,2]^. Mycetoma is endemic in many tropical and subtropical countries, these regions are referred to as the “mycetoma belt” that lies between 15°S – 30°N of the equator ^[3–6]^. Sudan is considered as one of the most affected regions by this disease, furthermore 70% of the prevalence causes of mycetoma is related to *Madurella mycetomatis* ^[6,2,5]^. The disease characterized by chronic subcutaneous masses, large tumor-like swellings, multiple draining sinuses, discharge grains, blood and pus; while the recurrent infection may lead to amputation ^[7,8,9,10,11]^. The infection is mainly located in the lower extremities, but could also affect other parts of the body ^[5,6]^. Mycetoma can affect all age groups (20-50 years) and is more common in males than females with a ratio 3:1. It is widely spread within people who had lived in rural areas and had worked in the farms due to their frequent contact to/with organisms, which have saprobiotic life in soil. In other hand, man to man infection does not occur ^[4]^. There are no ideal diagnostic tools available to identify mycetoma and the identification of the etiological agent is a very challenging issue that reflects directly on the treatment, especially in non-endemic areas ^[9,12]^. In the other hand treatment is not affordable, not effective, toxic for patients and requires a long period (18–24 months or more) ^[8,9]^. No vaccine was prepared yet against mycetoma therefore, vaccination is highly recommended ^[13]^. In a study done by Nele de Klerk et al, presents two antigenic proteins that have the ability to induce an antibody response in human fructose-bisphosphate aldolase (FBA) and pyruvate kinase (PK) of *M. mycetomatis* that might be useful as vaccine-candidates in the prevention of mycetoma ^[10]^.

The aim of the study is to predict effective epitope-based vaccine against fructose-bisphosphate aldolase (FBA) enzymes of *M. mycetomatis*. Development in immunogenetics will enhance apprehension of the impact of genetic factors on the interindividual and interpopulation variations in immune responses to vaccines that could be helpful for progress new vaccine strategies ^[14]^. *In silico* / reverse vaccinology had replaced conventional-culture based vaccine because it reduces the cost required for laboratory investigation of pathogen, also speeds up the time needed to achieve the results ^[15]^. Therefore, using immunoinformatics approaches to predict this new kind of vaccines could be a magnificently additive in the way forward of preventing mycetoma. Normally, the investigation of the binding affinity of antigenic peptides to the MHC molecules is the main goal when predicting epitopes. Using such tools and information leads to the development of new vaccines ^[15,16]^. While these approaches permit the optimization of a vaccine for a specific population, the problem can also be reformulated to design a “universal vaccine”: a vaccine that provides maximum coverage on the whole worlds’ population ^[17]^. In this study we focused on both MHC class II & I with performing of molecular docked in HLA-A0201.

## Materials and Methods

The Sequence of Fructose-bisphosphate aldolase (FBA) was retrieved from NCBI Database (https://www.ncbi.nlm.nih.gov/protein) ^[18]^ in a FASTA format as of September 2017 for further analysis, then the candidate epitopes were analyzed using different prediction tools of Immune Epitope Database IEDB analysis resource (http://www.iedb.org/) ^[19]^.

### 2.1. B-cell epitope prediction

Candidate epitopes were analyzed by several B-cell prediction methods that determine the antigenicity, hydrophilicity, flexibility and surface accessibility. The linear predicted epitopes were obtained by using BepiPred test from immune epitope database (http://tools.iedb.org/bcell/result/) ^[20]^ with a threshold value of 0.149 and a window size 6.

Furthermore, surface accessible epitopes were predicted with a threshold value of 1.0 and a window size of 6.0 using the Emini surface accessibility prediction tool ^[21]^.

The Antigenicity methods of Kolaskar and Tongaonker (http://tools.iedb.org/bcell/result/) were proposed to determine the sites of antigenic epitopes with a default threshold value of 1.030 and a window size of 6.0 ^[22]^.

### 2.2. MHC class I binding predictions

Analysis of peptide binding to MHC1 molecules was assessed by the IEDB MHC I prediction tool at http://tools.iedb.org/mhc1. The attachment of cleaved peptides to MHC molecules was predicted using artificial neural network (ANN) method. MHC1 alleles at score equal or less than 100 half-maximal inhibitory concentrations (IC50) were selected for further analysis ^[23,24]^.

### 2.3. MHC class II binding predictions

Analysis of peptide binding to MHC II molecules was assessed by the IEDB MHC II prediction tool at (http://tools.iedb.org/mhcii/result/). ^[25,26,27]^ For MHC II binding prediction, human allele references set were used. MHC II groove has the ability to bind different lengths peptides that makes prediction more difficult and less accurate. We used artificial neural networks to identify both the binding affinity and MHC II binding core epitopes. All epitopes that bind to many alleles at score equal or less than 1000 half-maximal inhibitory concentration (IC50) was selected for further analysis.

### 2.4. The Physicochemical properties

The physicochemical properties of Fructose-bisphosphate aldolase FBA protein was assessed using BioEdit sequence alignment editor software Version 7.2.5 ^[28]^.

### 2.5. Homology modeling

The reference sequence of Fructose-bisphosphate aldolase FBA) of *Madurella mycetomatis* to Raptor X web portal (http://raptorx.uchicago.edu) ^[29]^ in 19.09.2017, the 3D structure of it was received in 20.09./2017 and then treated with UCSF Chimera version 1.10.2, <u>UCSF ChimeraX</u> version 0.1 software to show the position of proposed peptides ^[30,31]^.

### 2.6. Protein structure retrieval and preparation

The HLA-A0201 was selected for docking and the 3-D structure of 4UQ3 was retrieved from the RCSB Protein Data Bank (http://www.rcsb.org/pdb/home/home.do). The structure 4UQ3 of this database was the crystal structure of HLA-A0201 in complex with an azobenzene-containing peptide ^[32]^. The protein files were prepared by removal of all water molecules and hetero groups.

### 2.7. *In silico* molecular docking

Molecular docking was performed using Autodock 4.0 software ^[33]^, based on Lamarckian Genetic Algorithm; which combines energy evaluation through grids of affinity potential to find the suitable binding position for a ligand on a given protein ^[34]^. Polar hydrogen atoms were added to the protein targets and Kollman united atomic charges were computed. All hydrogen atoms were added to the ligands before the Gastiger partial charges were assigned, then a co-crystal ligand was removed and the bond orders were checked. The target’s grid map was calculated and set to 60×60×60 points with grid spacing of 0.375 Å. The grid box was then allocated properly in the target to include the active residue in the center. The default docking algorithms were set in accordance with standard docking protocol. Finally, ten independent docking runs were carried out for each ligand Peptide and results were retrieved as binding energies. Poses that showed lowest binding energies were visualized using UCSF chimera ^[35]^

## 3. Results

### 3.1. B-cell epitope prediction

The sequence of Fructose-bisphosphate aldolase [Madurella mycetomatis] was subjected to Bepipred linear epitope prediction, Emini surface accessibility and Kolaskar and Tongaonkar antigenicity methods in I DB, to determine the binding to B cell, being in the surface and to test the immunogenicity. The results were shown in Table (1) & Figure (1-4).

**Table (1):**
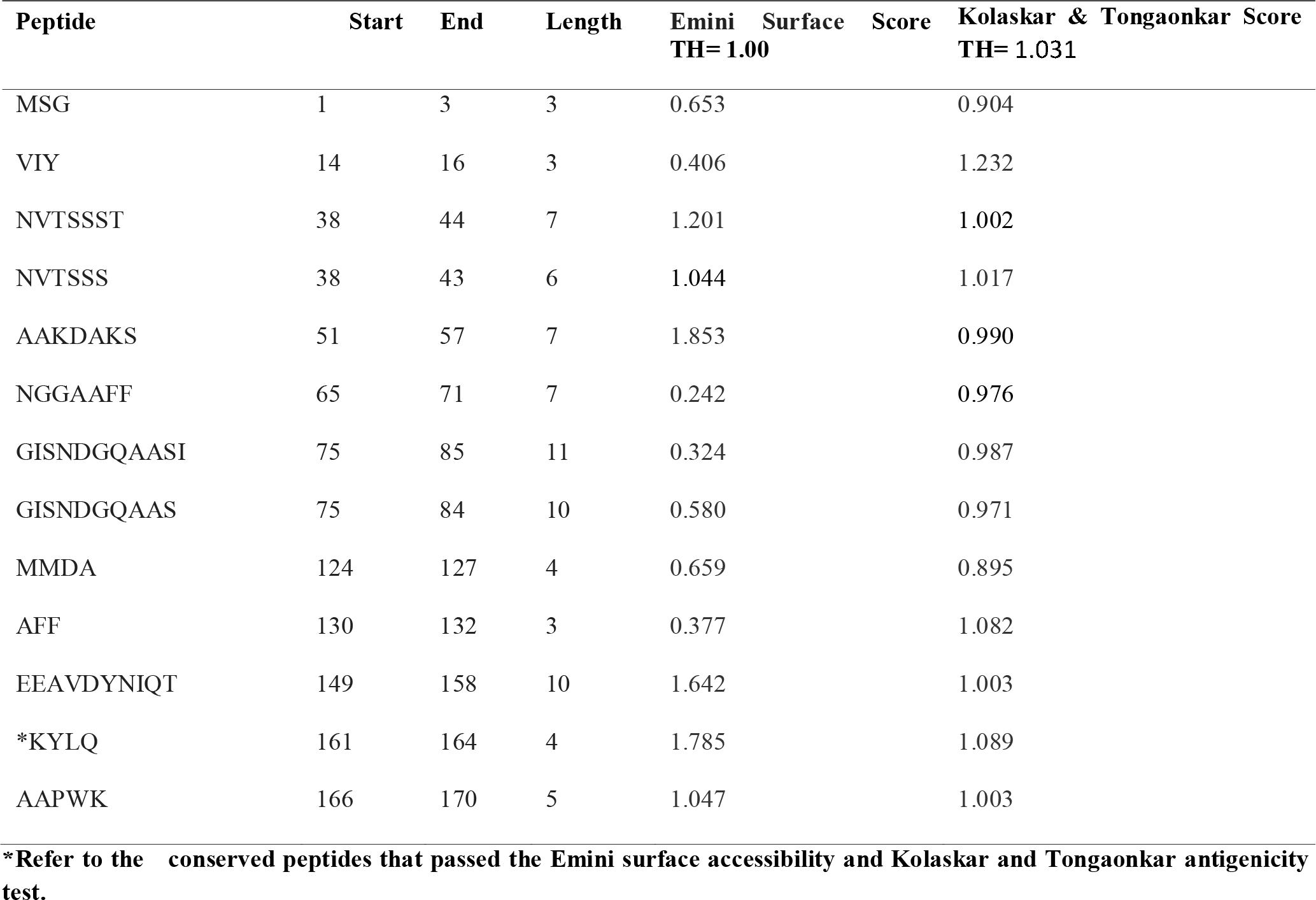
List of peptides with their surface accessibility score and antigenicity score.

**Figure (1).**
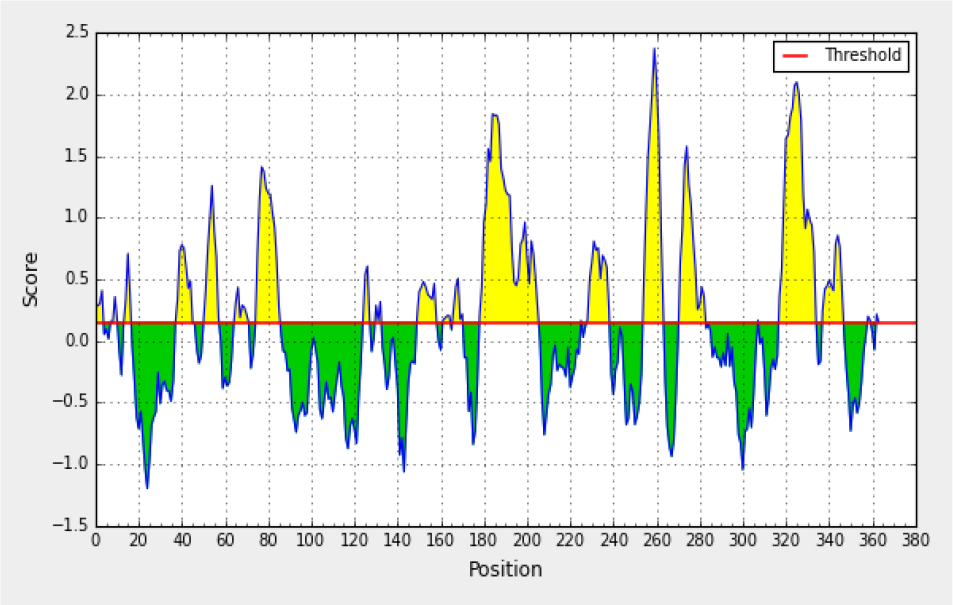
Present <u>Bepipred Linear Epitope Prediction</u>, the yellow space above threshold (red line) is proposed to be a part of B cellepitopes and

**Figure (2).**
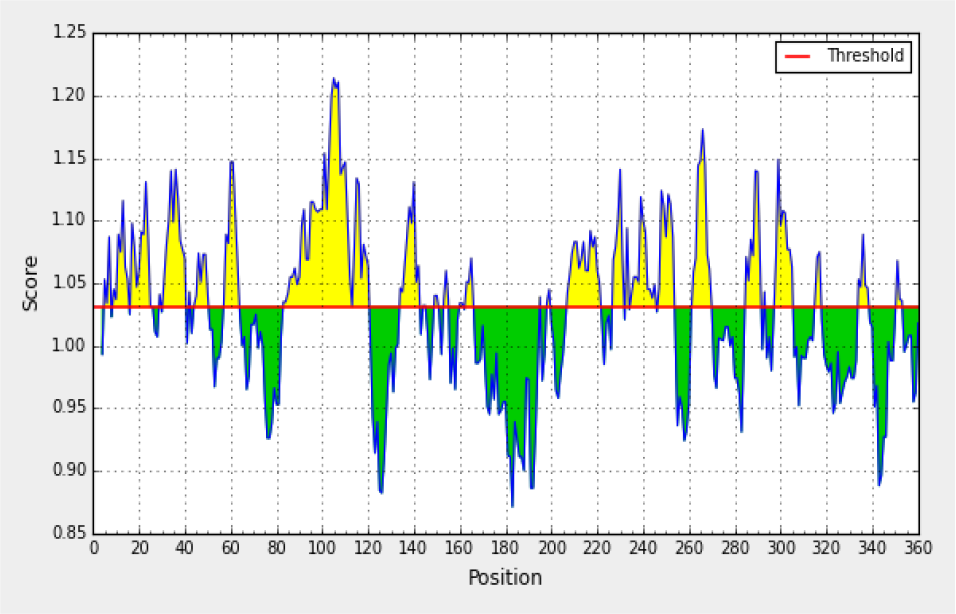
Present Kolaskar and Tongaonkar antigenicity prediction, Yellow areas above threshold (red line) are proposed to be a part of B cell epitope

**Figure (3).**
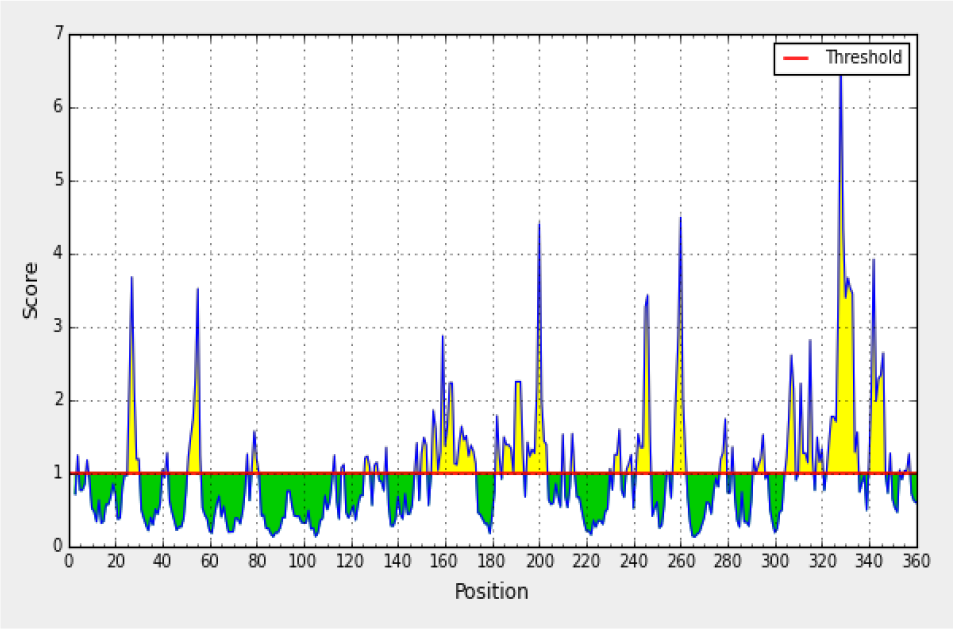
Present Emini surface accessibility prediction, the yellow space above threshold (red line) is proposed to be a part of B cell epitopes and the green space is not a part.

**Figure (4).**
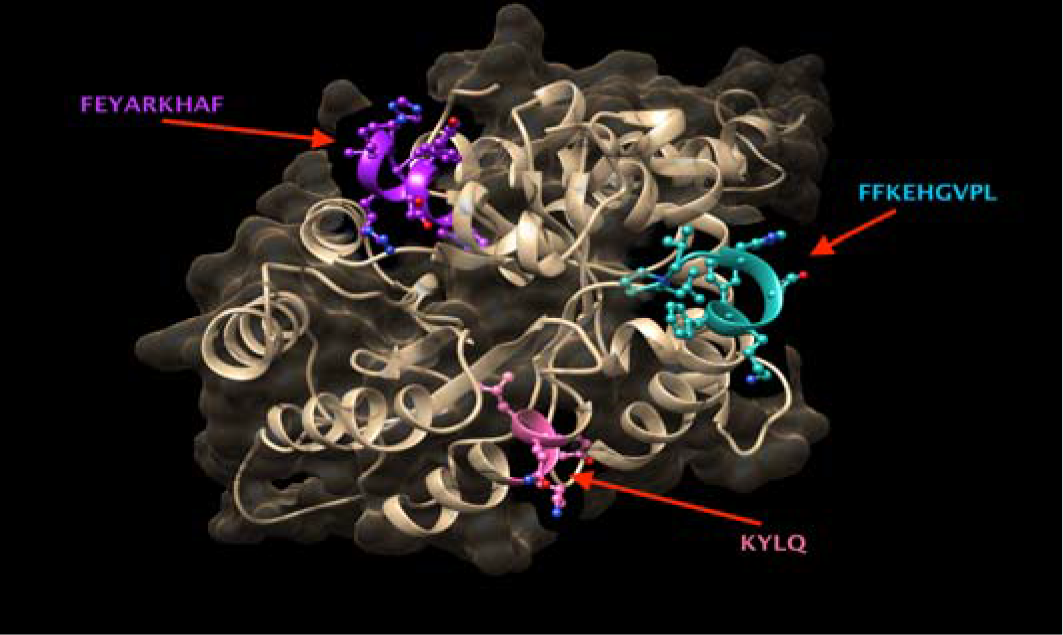
The Structural position of the most promising B-cell epitope peptide KYLQ, FEYARKHAF of MHC I and FFKEHGVPL MHC I and MHC II of Fructose-bisphosphate aldolase [Madurella mycetomatis] using UCSF Chimera version 1.10.2.

### 3.2. Prediction of T helper cell epitopes and interaction with MHC I alleles

Fructose-bisphosphate aldolase [Madurella mycetomatis] sequence was analyzed using IEDB MHC-I binding prediction tool based on ANN-align with half-maximal inhibitory concentration (IC_50_) ≤100; the list most promising epitopes that had Binding affinity with the Class I alleles along with their positions in the Fructose-bisphosphate aldolase [Madurella mycetomatis] were shown in table (2).

**Table (2):**
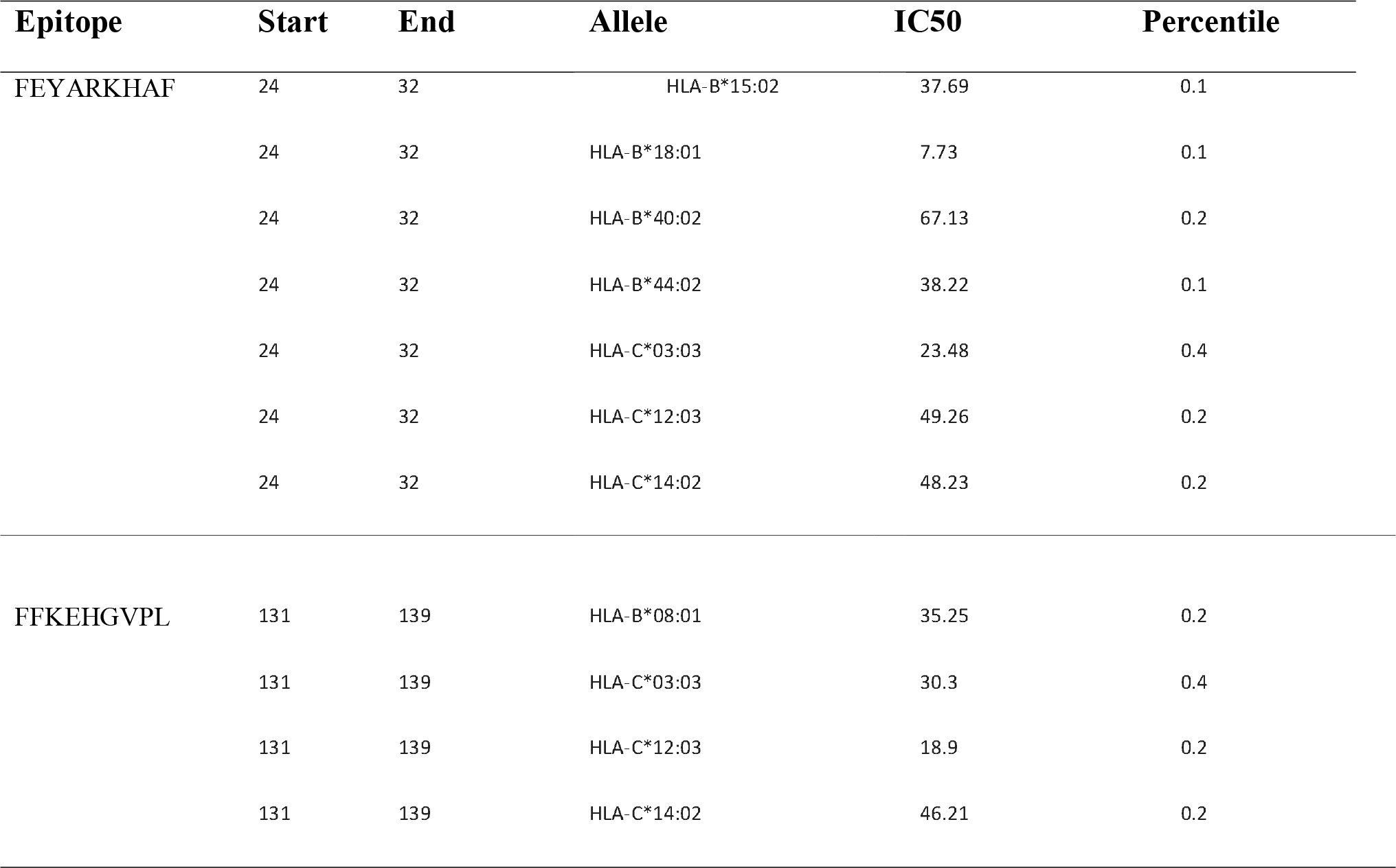
List of most promising epitopes that had Binding affinity with MHC-I alleles along with their positions in the Fructose-bisphosphate aldolase [Madurella mycetomatis], IC50 and Percentile.

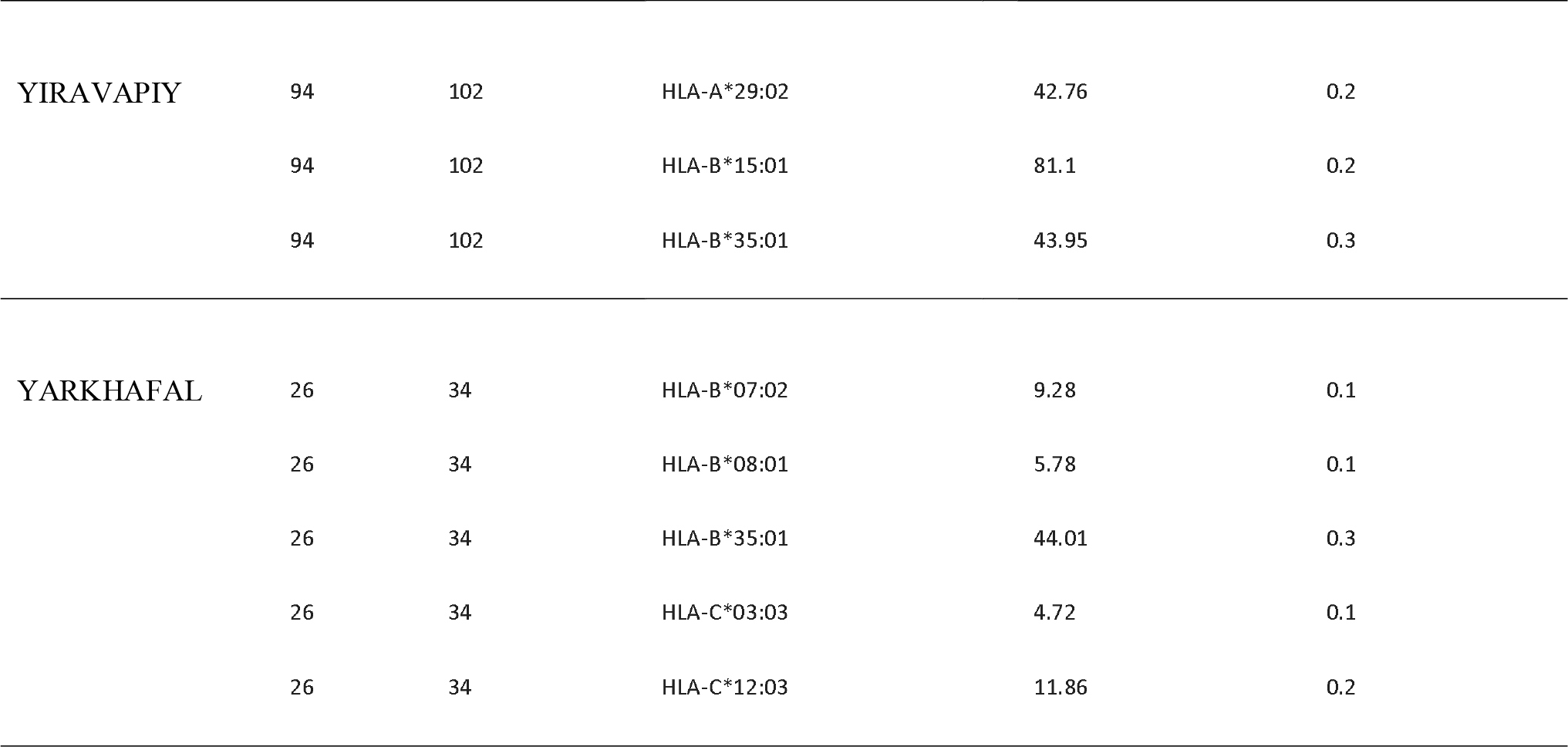

### 3.3. Prediction of T helper cell epitopes and interaction with MHC II alleles

Fructose-bisphosphate aldolase [Madurella mycetomatis] sequence was analyzed using IEDB MHC-II binding prediction tool based on NN-align with half-maximal inhibitory concentration (IC_50_) ≤1000; the list of the most promising epitopes and their correspondent binding MHC11 alleles along with their positions in the Fructose-bisphosphate aldolase [Madurella mycetomatis] were shown in (supplementary table 1) while the list most promising epitopes that had a strongly Binding affinity with the Class II alleles and the number of their binding alleles were shown in table (3).

**Table (3):**
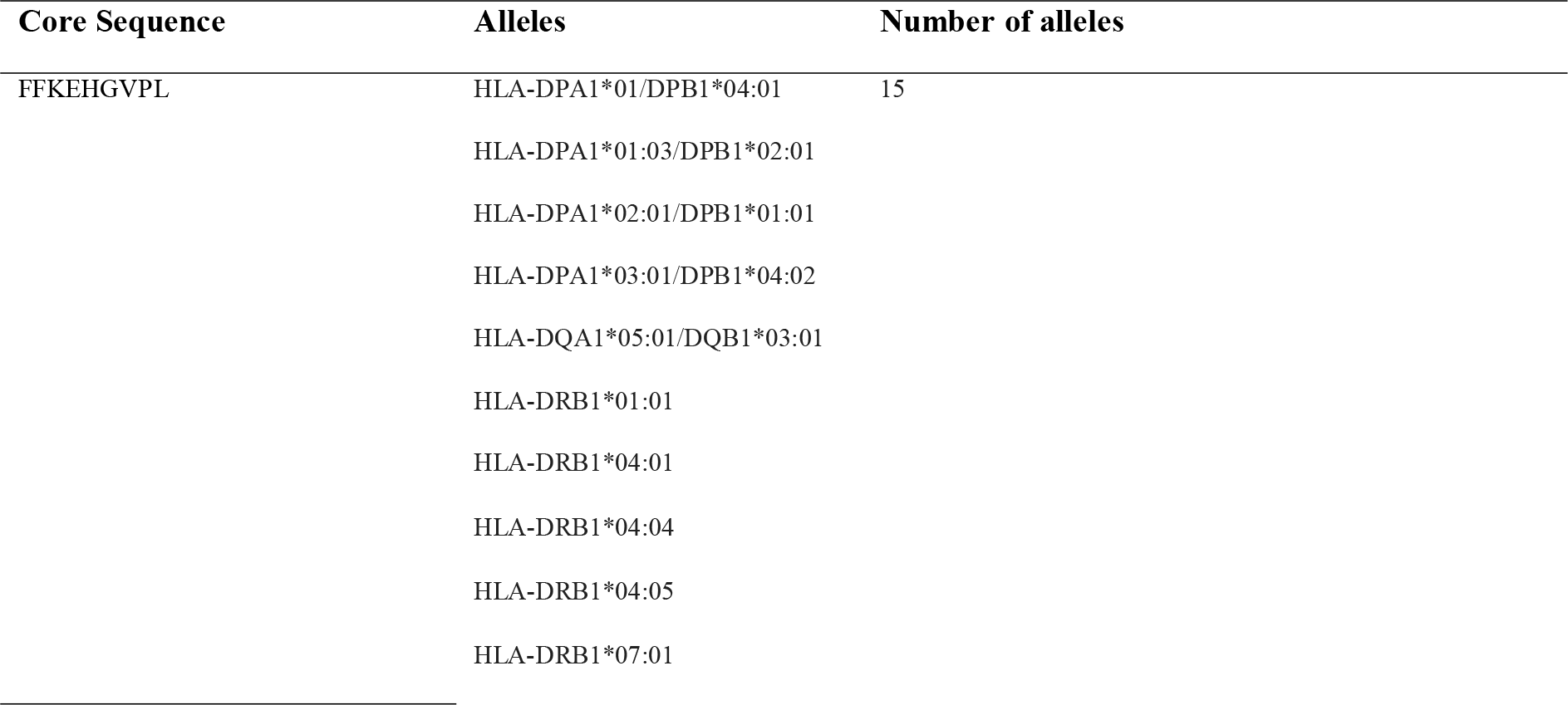
List of most promising epitopes (Core Sequence) that had Binding affinity with MHC-II alleles and the number of their binding alleles.

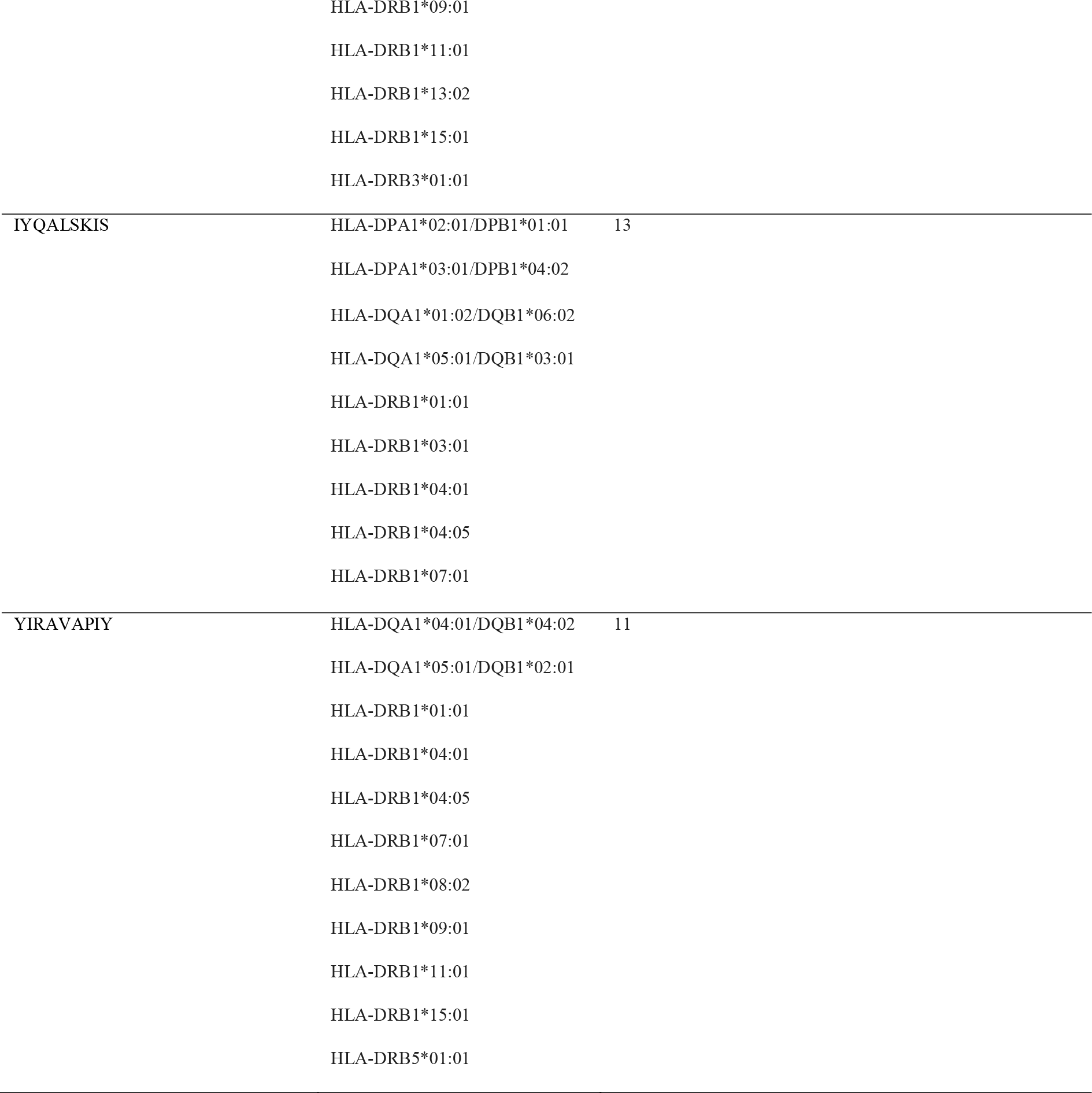

### 3.4. Fructose-bisphosphate aldolase physical and chemical parameters

The physicochemical properties of Fructose-bisphosphate aldolase (FBA) protein was assessed using BioEdit software version 7.0.9.0. The protein length was found to be 363 amino acids and the molecular weight was 39675.83 Daltons. The amino acid that formed FBA protein and their number along with their molar percentage (Mol%) were shown in table (4).

**Table (4):** List of amino acid that formed Fructose-bisphosphate aldolase (FBA) protein, their number and Mol% using BioEdit software Version 7.2.5.

| Amino Acid | Number | Mol% | Amino Acid | Number | Mol% |
| --- | --- | --- | --- | --- | --- |
| Ala A | 40 | 11.02 | Met M | 7 | 1.93 |
| Cys C | 3 | 0.83 | Asn N | 16 | 4.41 |
| Asp D | 26 | 7.16 | Pro P | 17 | 4.68 |
| Glu E | 18 | 4.96 | Gln Q | 13 | 3.58 |
| Phe F | 16 | 4.41 | Arg R | 9 | 2.48 |
| Gly G | 32 | 8.82 | Ser S | 19 | 5.23 |
| His H | 12 | 3.31 | Thr T | 14 | 3.86 |
| Ile I | 20 | 5.51 | Val V | 28 | 7.71 |
| Lys K | 26 | 7.16 | Trp W | 6 | 1.65 |
| Leu L | 27 | 7.44 | Tyr Y | 14 | 3.86 |

**Figure (5).**
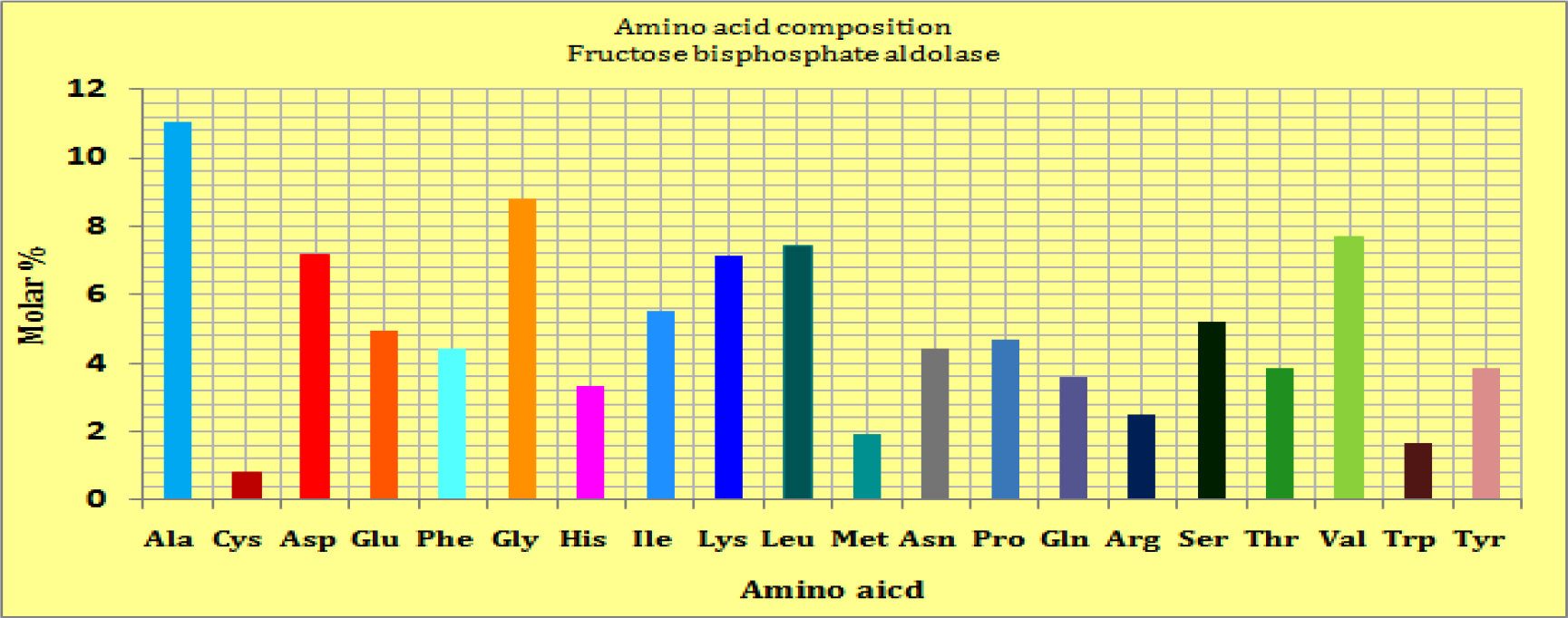
The graph shows the Amino Acid Composition Mol% of Fructose bisphosphate aldolase using Bio Edit software Version 7.2.5.

### 3.5. Population Coverage Analysis

Population coverage test was performed to detect all epitopes binds to MHC1 alleles, MHC11 alleles for Sudan and North Africa.

**Table (5):** Population coverage average for all epitopes binding to MHC1 alleles in World.

| Population/area | Class I |  |  |
| --- | --- | --- | --- |
|  | Coverage<sup>a</sup> | Average hit<sup>b</sup> | pc90<sup>c</sup> |
| **<u>World</u>** | 97.57% | 8.77 | 3.22 |
| **Average** | 97.57 | 8.77 | 3.22 |
| **Standard deviation** | 0.0 | 0.0 | 0.0 |

<sup>a</sup> Projected population coverage
<sup>b</sup> Average number of epitope hits / HLA combinations recognized by the population
<sup>c</sup> Minimum number of epitope hits / HLA combinations recognized by 90% of the population

**Table (6):** Population coverage average for all epitopes binding to MHC1 alleles in North Africa.

| Population/area | Class I |  |  |
| --- | --- | --- | --- |
|  | Coverage<sup>a</sup> | Average hit<sup>b</sup> | pc90<sup>c</sup> |
| **<u>North Africa</u>** | 94.16% | 7.35 | 1.91 |
| **Average** | 94.16 | 7.35 | 1.91 |
| **Standard deviation** | 0.0 | 0.0 | 0.0 |

<sup>a</sup> Projected population coverage
<sup>b</sup> Average number of epitope hits / HLA combinations recognized by the population
<sup>c</sup> Minimum number of epitope hits / HLA combinations recognized by 90% of the population

**Table (7):** Population coverage average for all epitopes binding to MHC1 alleles in Sudan.

| Population/area | Class I |  |  |
| --- | --- | --- | --- |
|  | Coverage<sup>a</sup> | Average hit<sup>b</sup> | pc90<sup>c</sup> |
| **<u>Sudan</u>** | 92.64% | 7.12 | 1.48 |
| **Average** | 92.64 | 7.12 | 1.48 |
| **Standard deviation** | 0.0 | 0.0 | 0.0 |

<sup>a</sup> Projected population coverage
<sup>b</sup> Average number of epitope hits / HLA combinations recognized by the population
<sup>c</sup> Minimum number of epitope hits / HLA combinations recognized by 90% of the population

**Table (8):** Population coverage average for all epitopes binding to MHC11 alleles in World.

| Population/area | Class II |  |  |
| --- | --- | --- | --- |
|  | Coverage<sup>a</sup> | Average hit<sup>b</sup> | pc90<sup>c</sup> |
| **<u>World</u>** | 81.94% | 41.15 | 8.86 |
| **Average** | 81.94 | 41.15 | 8.86 |
| **Standard deviation** | 0.0 | 0.0 | 0.0 |

<sup>a</sup> Projected population coverage
<sup>b</sup> Average number of epitope hits / HLA combinations recognized by the population
<sup>c</sup> Minimum number of epitope hits / HLA combinations recognized by 90% of the population

**Table (9):** Population coverage average for all epitopes binding to MHC11 alleles in North Africa.

| Population/area | Class II |  |  |
| --- | --- | --- | --- |
|  | <sup>a</sup><br>Coverage | <sup>b</sup><br>Average hit | <sup>c</sup><br>pc90 |
| **<u>North Africa</u>** | 75.2% | 30.7 | 6.45 |
| **Average** | 75.2% | 30.7 | 6.45 |
| **Standard deviation** | 0.0 | 0.0 | 0.0 |

<sup>a</sup> Projected population coverage
<sup>b</sup> Average number of epitope hits / HLA combinations recognized by the population
<sup>c</sup> Minimum number of epitope hits / HLA combinations recognized by 90% of the population

**Table (10):** Population coverage average for all epitopes binding to MHC11 alleles in Sudan.

| Population/area | Class II |  |  |
| --- | --- | --- | --- |
|  | <sup>a</sup><br>Coverage | <sup>b</sup><br>Average hit | <sup>c</sup><br>pc90 |
| **<u>Sudan</u>** | 59.3% | 20.73 | 3.93 |
| **Average** | 59.3 | 20.73 | 3.93 |
| **Standard deviation** | 0.0 | 0.0 | 0.0 |

<sup>a</sup> Projected population coverage
<sup>b</sup> Average number of epitope hits / HLA combinations recognized by the population
<sup>c</sup> Minimum number of epitope hits / HLA combinations recognized by 90% of the population

### 3.6. Molecular Docking

**Figure (6).**
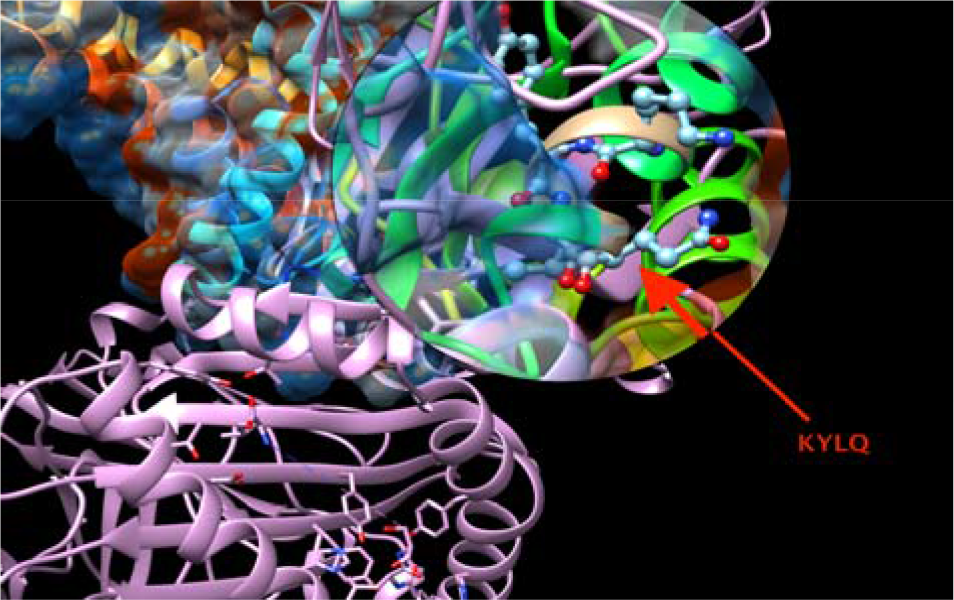
Molecular Docking of KYLQ peptide of Fructose-bisphosphate aldolase [Madurella mycetomatis] that binds B-cell and docked in HLA-A0201 using UCSF Chimera version 1.10.2.

**Figure (7).**
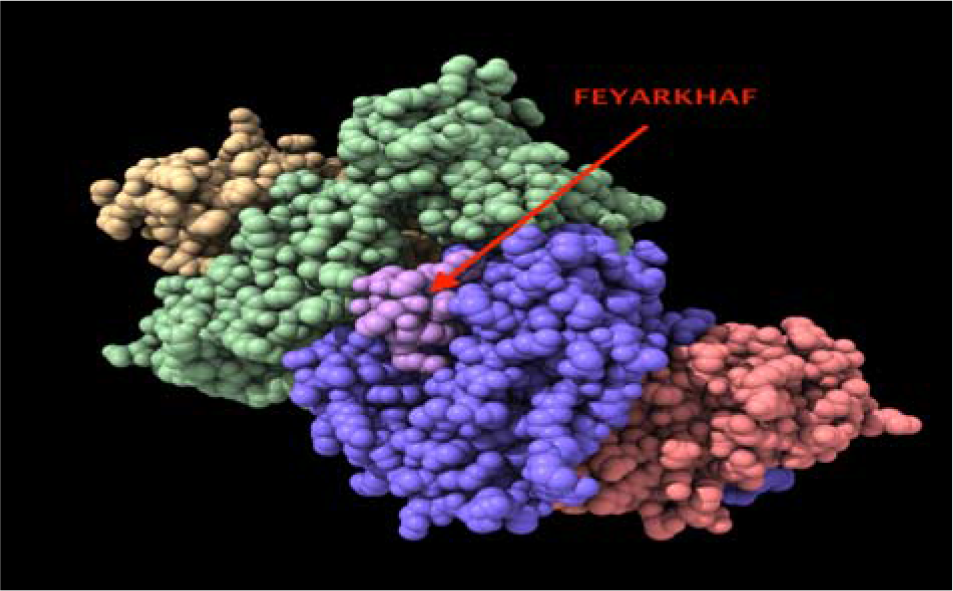
Molecular Docking of the most promising peptide FEYARKHAF of Fructose-bisphosphate aldolase [Madurella mycetomatis] that binds MHC I alleles and docked in HLA-A0201 using UCSF Chimera X version 0.1

**Figure (8).**
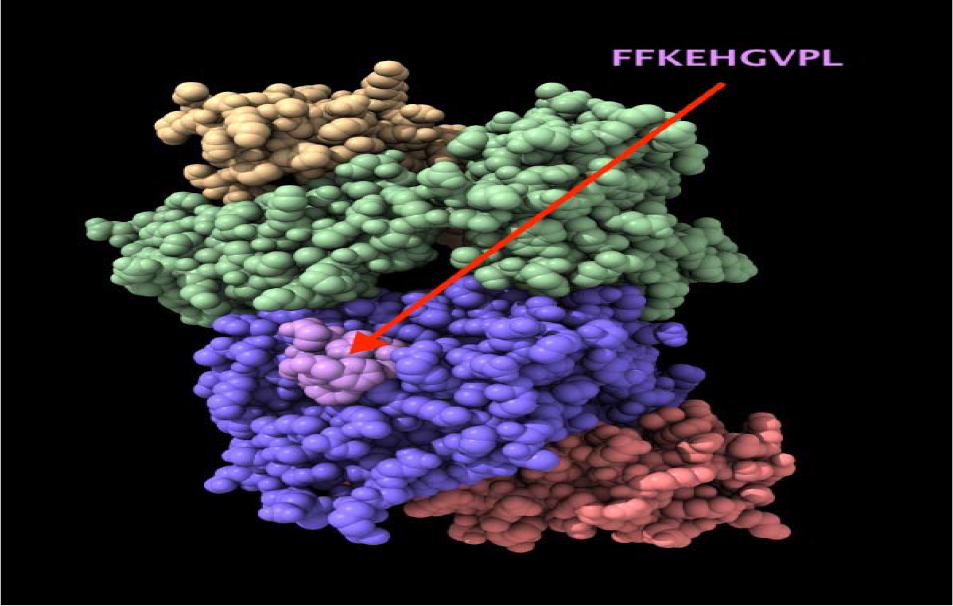
Molecular Docking of the most promising peptide FFKEHGVPL of Fructose-bisphosphate aldolase [Madurella mycetomatis] that binds MHC II alleles and MHC I and docked in HLA-A0201 using UCSF Chimera X version 0.1

## Discussion

The current study reveled three promising peptides that could be the first vaccine against Fructose bisphosphate aldolase *madurella mycetomatsis* worldwide. The peptide KYLQ is the only peptide that passed all B-cell prediction tests which used in this study and show high binding affinity in docking. In MHC 71 conserved peptides were predicted using (ANN) method with (IC50) ≤ 100. 3 peptides were suggested and thought to be a possible epitope based peptide vaccine against *madurella mycetomatsis*. The peptide FEYARKHAF interact with the highest number of alleles and the other two peptides YIRAVAPIY and FFKEHGVPL are highly binding peptides. The selection was done according to the binding with the highest numbers of alleles. FEYARKHAF interact with 7alleles(HLA-B*15:02,HLA-B*18:01,HLA-B*40:02,HLA-B*44:02,HLA-C*03:03,HLA-C*12:03,HLA-C*14:02), YIRAVAPIY that binds with 3Alleles (HLA-A*29:02,HLA-B*15:01,HLA-B*35:01) and FFKEHGVPL that interact with 4 alleles (HLA-B*08:01,HLA-C*03:03,HLA-C*12:03,HLA-C*14:02). The reference FBA was analyzed using IEDB MHC II binding prediction tool based on NN-align with half-maximal inhibitory concentration (IC50) ≤ 1000; there were 240 predicted peptides found to interact with MHC-II alleles. We proposed the peptides FFKEHGVPL, IYQALSKIS and YIRAVAPIY that had the affinity to bind the highest number of MHC11 alleles. The peptide FFKEHGVPL exhibits exceptional population coverage results for both MHC1 and MHC11 alleles with total binding to 15 different alleles in MHC 11 and four alleles in MHC1, This finding show a very strong potential to formulate an epitopes-based peptide vaccine against *madurella mycetomatsis* and make the peptide FFKEHGVPL highly proposed. The Population Coverage analysis using IEDB analysis resource predicted both MHC class I and class II based coverage of the selected peptides for Sudan, North Africa and world population to assess the feasibility of being potential vaccine candidate in those endemic aeries. The peptide FEYARKHAF population coverage was found to be 30.3% for Sudan, 24.38% for North Africa and the world population coverage was found to be 36.63%. Furthermore the peptide FFKEHGVPL population coverage was found to be 26.59%for Sudan, 20.37%for North Africa and the world population coverage was found to be 29.11%.

Population coverage results for total peptides binding to MHC1 alleles with IC50≤100 shows 97.57% projected population coverage for the world, 94.16% in North Africa and 92.64% in Sudan while the population coverage results for total peptides binding to MHC II alleles Shows 81.94% projected population coverage for the world, 75.2% in North Africa and 59.3%in Sudan. Finally, three peptides KYLQ, FEYARKHAF and FFKEHGVPL were proposed as potential candidates to be devolved in to a vaccine. Herein, *in-silico* molecular docking was done to explore the binding affinity between the aforementioned peptides and the target HLA-A0201. This target has been selected for docking pertained with its involvement in several immunological and pathological diseases hence, although numerous studies have demonstrated an association between HLA alleles and disease susceptibility, Defining protective HLA allelic associations potentially allows the identification of pathogen epitopes that are restricted by the specific HLA alleles. These epitopes may then be incorporated into vaccine design in the expectation that the natural resistance can be replicated by immunization ^[36,37,38]^. The binding energies between the peptides and HLA-A0201 molecule were calculated and ranked. Peptide FFKEHGVPL shows the best Docking result, this indicates a strong potential to formulate a vaccine utilizing the peptide FFKEHGVPL that is highly promising to be the first proposed epitope-based peptide vaccines against Fructose-bisphosphate aldolase (FBA) of *Madurella mycetomatis*. Based on our previous studies on Mokola Rabies Virus and Lagos Rabies Virus, a computational analysis was made and the most immunogenic epitopes for T and B cells involved in the cell-mediated immunity were analyzed ^[39–40]^. In this study the same techniques were used, to design the Epitope-Based Peptide Vaccine against fructose-bisphosphate aldolase (FBA) enzymes of *M. mycetomatis* as an immunogenic part to stimulate a protective immune response.

One of the limitations of this study was the number of Fructose-bisphosphate aldolase (FBA) Sequences; there was only one Sequence of FBA that retrieved from NCBI Database: This Sequence collected from Sudan. On the other hand, the new version of IEDB population coverage calculation tool does not illustrate the ability to know the Population coverage results for every peptide binding to MHC11 like as previously was, it illustrates only the average percentage of all peptides. Recently, two other studies were going along parallel with our study to predict new epitope-based peptide vaccines against *Madurella mycetomatis*. They used Pyruvate Kinase (PK) of *Madurella mycetomatis* and Translationaly Controlled Tumor Protein (TCTP) as target proteins ^[41,42]^. But we believe that fructose-bisphosphate aldolase (FBA) enzymes could be the best target protein due to its high ability to induce an antibody response in a human. In Nele de Klerk et al study, they found that both FBA and PK IgG antibodies were present in eumycetoma patients’ sera. However, only FBA antibody levels were found to be significantly higher in eumycetoma patients’ sera when compared to healthy controls ^[10]^.

## Conclusion

To the best of our knowledge this study is consider to be the first to propose epitope-based peptide vaccine against Fructose-bisphosphate aldolase (FBA) of *Madurella mycetomatis*, which is expected to be highly antigenic with a minimum allergic effect. Furthermore, this study proposes promising peptides of (KYLQ) that show a very strong binding affinity to B-cell, (FEYARKHAF) with a very strong binding affinity to (MHC1) alleles and (FFKEHGVPL) that shows a very strong binding affinity to (MHC11) and (MHC1) alleles. In-vivo and in-vitro assessment for the most promising peptides namely; (KYLQ), (FEYARKHAF) and (FFKEHGVPL) are recommended to be explored and find out more on their ability to be developed into vaccines against *Madurella mycetomatis*.

## Competing interests

The authors declare that they have no competing interests.

## References

1. Zijlstra EE, van de Sande WWJ, Welsh O, Mahgoub ES heikh, Goodfellow M, Fahal, AH. Mycetoma: a unique neglected tropical disease. Lancet Infect Dis [Internet]. Elsevier Ltd; 2016;16(1):100–12. Available from: http://dx.doi.org/10.1016/S1473-3099(15)00359-X.

2. Fahal A, Mahgoub ES, Hassan AMEL, Abdel-Rahman ME. Mycetoma in the Sudan: An Update from the Mycetoma Research Centre, University of Khartoum, Sudan. PLoS Negl Trop Dis. 2015;9(3):1–19.

3. Samy AM, van de Sande WWJ, Fahal AH, Peterson, AT. Mapping the potential risk of mycetoma infection in Sudan and South Sudan using ecological niche modeling. PLoS Negl Trop Dis [Internet]. 2014;8(10):e3250.Availablefrom: http://www.pubmedcentral.nih.gov/articlerender.fcgi?artid=4199553&tool=pmcentrez&rendertype=abstract

4. Efared B, Tahiri L, Boubacar MS, Atsam-Ebang G, Hammas N, Hinde EF, et al. Mycetoma in a non-endemic area: a diagnostic challenge. BMC Clin Pathol [Internet]. BMC Clinical Pathology; 2017;17(1):1. Available from: http://bmcclinpathol.biomedcentral.com/articles/10.1186/s12907-017-0040-5

5. Fahal AH, Van De Sande WWJ. The Epidemiology of Mycetoma. 2012;320–6.

6. Brufman T, Ben-Ami R, Mizrahi M, Bash E, Paran, Y. Mycetoma of the foot caused by madurella mycetomatis in immigrants from sudan. Isr Med Assoc J. 2015;17(7):418–20.

7. Elhassan M, Yousif A, Elmekki M, Hamid, M. Isolation and molecular identification of actinomycetes from mycetoma patients in Sudan. Ann Trop Med Public Heal. 2013;6(2):211–4.

8. Van de Sande WWJ, Maghoub ES, Fahal AH, Goodfellow M, Welsh O, Zijlstra, EE. The Mycetoma Knowledge Gap: Identification of Research Priorities. PLoS Negl Trop Dis. 2014;8(3): e2667. doi:10.1371/journal.pntd.0002667.

9. Van de Sande WWJ, Fahal AH, Goodfellow M, Mahgoub ES, Welsh O, Zijlstra, EE. Merits and Pitfalls of Currently Used Diagnostic Tools in Mycetoma. PLoS Negl Trop Dis. 2014; 8(7): e2918. http://doi.org/10.1371/journal.pntd.0002918.

10. Nele de Klerk, Corné de Vogel, Ahmed Fahal, Alex van Belkum, Wendy, W. J. van de Sande; Fructose-bisphosphate aldolase and pyruvate kinase, two novel immunogens in Madurella mycetomatis, Medical Mycology, Volume 50, Issue 2, 1 February 2012, Pages 143–151, http://doi.org/10.3109/13693786.2011.593005.

11. Van de Sande WWJ, Janse DJ, Hira V, Goedhart H, van der Zee R, Ahmed AO, et al. Translationally Controlled Tumor Protein from Madurella mycetomatis: a Marker for Tumorous Mycetoma Progression. The Jour of Imm. 2006;177: 1997–2005.

12. Van de Sande WW, Maghoub el S, Fahal AH, Goodfellow M, Welsh O, Zijlstra E. The mycetoma knowledge gap: identification of research priorities. PLoS Negl Trop Dis. 2014;8(3): e2667.

13. Hay RJ, Fahal, AH. Mycetoma: an old and still neglected tropical disease, Trans R Soc Trop Med Hyg. 2015;109(3):169–170. http://doi.org/10.1093/trstmh/trv003.

14. Lemaire D, Barbosa T, Rihet, P. Coping with genetic diversity: the contribution of pathogen and human genomics to modern vaccinology. Braz J Med Biol Res. 2012;45:376–385. Available from: http://www.ncbi.nlm.nih.gov/pmc/articles/PMC3854287/ doi: 10.1590/S0100-879X2011007500142.

15. Patronov A, Doytchinova, I. T-cell epitope vaccine design by immunoinformatics. Open Biol. 2013;3(1):120139. doi:10.1098/rsob.120139.

16. Tomar N, De, RK. 2010 Immunoinformatics: an integrated scenario. Immunology 113, 153–168. (doi:10.1111/j.1365-2567.2010.03330.x).

17. Toussaint NC, Maman Y, Kohlbacher O, Louzoun, Y. Universal peptide vaccines - optimal peptide vaccine design based on viral sequence conservation. Vaccine. 2011;29:8745–53. doi:10.1016/j.vaccine.2011.07.132.

18. <http://www.ncbi.nlm.nih.gov/protein>

19. Vita R, Overton JA, Greenbaum JA, Ponomarenko J, Clark JD, Cantrell JR, Wheeler DK, Gabbard JL, Hix D, Sette A, Peters, B. The immune epitope database (IEDB) 3.0. Nucleic Acids Res. 2014 Oct 9 pii: gku938. [Epub ahead of print] PubMed PMID: 25300482.

20. Larsen JE, Lund O, Nielsen M (2006) Improved method for predicting linear B-cell epitopes. Immunome Res 2: 2.

21. Emini EA, Hughes JV, Perlow D, Boger J (1985) Induction of hepatitis A virus-neutralizing antibody by a virus-specific synthetic peptide. Journal of virology 55: 836–839.

22. Kolaskar AS, Tongaonkar PC (1990) A semi-empirical method for prediction of antigenic determinants on protein antigens. FEBS Lett 276: 172–174.

23. Nielsen M, Lundegaard C, Worning P, Lauemoller SL, Lamberth K, et al. (2003) Reliable prediction of T-cell epitopes using neural networks with novel sequence representations. Protein Sci 12: 1007–1017.

24. Kim Y, Ponomarenko J, Zhu Z, Tamang D, Wang P, et al. (2012) Immune epitope database analysis resource. Nucleic Acids Res 40: W525–530.

25. Nielsen M, Lundegaard C, Worning P, Lauemoller SL, Lamberth K, et al. (2003) Reliable prediction of T-cell epitopes using neural networks with novel sequence representations. Protein Sci 12: 1007–1017.

26. Kim Y, Ponomarenko J, Zhu Z, Tamang D, Wang P, et al. (2012) Immune epitope database analysis resource. Nucleic Acids Res 40: W525–530.

27. Wang P, Sidney J, Dow C, Mothe B, Sette A (2008) A systematic assessment of MHC class II peptide binding predictions and evaluation of a consensus approach. PLoS Comput Biol 4: e1000048.

28. Hall TA (1999) BioEdit: a user-friendly biological sequence alignment editor and analysis program for Windows 95/98/NT. Nucl Acids Symp Ser 41: 95–98.

29. <http://raptorx.uchicago.edu/>

30. UCSF Chimera--a visualization system for exploratory research and analysis. Pettersen EF, Goddard TD, Huang CC, Couch GS, Greenblatt DM, Meng EC, Ferrin, TE. J Comput Chem. 2004 Oct;25(13):1605–12.

31. UCSF ChimeraX: Meeting Modern Challenges in Visualization and Analysis. Goddard TD, Huang CC, Meng EC, Pettersen EF, Couch GS, Morris JH, Ferrin, TE. Protein Sci. 2018 Jan;27(1):14–25. doi: 10.1002/pro.3235.

32. Choo, J.A.; Thong, S.Y.; Yap, J.; van Esch, W.J.; Raida, M.; Meijers, R.; Lescar, J.; Verhelst, S.H.; Grotenbreg, G.M. Bioorthogonal cleavage and exchange of major histocompatibility complex ligands by employing azobenzene-containing peptides. Angew. Chem. 2014, 53, 13390–13394.

33. Morris GM, Goodsell DS, Halliday RS, Huey R, Hart WE, Belew RK, Olson, AJ. Automated docking using a Lamarckian genetic algorithm and empirical binding free energy function. J Comput Chem, 1998;19:1639–62.

34. Morris GM, Huey R, Lindstrom W, Sanner M F, Belew RK, Goodsell DS, Olson, AJ. Autodock4 and AutoDockTools4: automated docking with selective receptor flexiblity, J Comput Chem, 2009;30:2785–91.

35. Pettersen EF, Goddard TD, Huang CC, Couch GS, Greenblatt DM, Meng EC, Ferrin, TE. UCSF Chimera-a visualization system for exploratory research and analysis. J Comput Chem, 2004;25:1605–12.

36. Burton, P. R., M. D. Tobin, and J. L. Hopper. 2005. Key concepts in genetic epidemiology. Lancet 366:941–951. [PubMed].

37. Calafell, F., and N. Malats. 2003. Basic molecular genetics for epidemiologists. J. Epidemiol. Community Health 57:398–400. [PMC free article] [PubMed].

38. Davenport, M. P., and A. V. Hill. 1996. Reverse immunogenetics: from HLA-disease associations to vaccine candidates. Mol. Med. Today. 2:38–45. [PubMed].

39. Mohammed AA, Hashim O, Elrahman KAA, Hamdi A, Hassan MA (2017) Epitope-Based Peptide Vaccine Design Against Mokola Rabies Virus Glycoprotein G Utilizing In Silico Approaches. Immunome Res 13: 144. Doi: 10.4172/1745-7580.1000144.

40. Ahmed OH, Abdelhalim A, Obi S, Abd_elrahman KA, Hamdi A, et al. (2017) Immunoinformatic Approach for Epitope-Based Peptide Vaccine against Lagos Rabies Virus Glycoprotein G. Immunome Res 13: 137. doi: 10.4172/17457580.1000137.

41. Aya Yusri A. Manofali, Ismail, M. A. I, Reem E. Talha, et al. (2018) Vaccinomics Approach for Designing Potential Peptide Vaccine by Targeting Pyruvate Kinase of Madurella Mycetomatis. (Unpublished).

42. Ahmed.O.H, Walaa.A.Omer, Samira.M.Polis, et al. (2018) Immunoinformatics Prediction of Epitope Based Peptide Vaccine Against Madurella mycetomatis Translationally Controlled Tumor Protein (Unpublished).

